# mim: A lightweight auxiliary index to enable fast, parallel, gzipped FASTQ parsing

**DOI:** 10.1101/2025.11.24.690271

**Authors:** Rob Patro, Siddhant Bharti, Prajwal Singhania, Rakrish Dhakal, Thomas J. Dahlstrom, Ragnar Groot Koerkamp

**Affiliations:** Department of Computer Science, University of Maryland, College Park, MD 20742, United States; Center for Bioinformatics and Computational Biology, University of Maryland

**Keywords:** `FASTQ`, parsing, parallel, compression, genomics, sequencing

## Abstract

The `FASTQ` file format is the *lingua franca* of primary data distribution and processing across most of bioinformatics. Over time, the compression, storage, transmission, and decompression of `gzip` compressed `fastq.gz` files has become a substantial scalability bottleneck in the modern world of fast and massively parallel genomics tools and algorithms.

In this work, we introduce `mim`: a lightweight, *auxiliary* index that enables fast, parallel, and highly-scalable parsing of compressed `fastq.gz` files. The creation of the `mim` index for a file is a one-time operation that can be performed in time comparable to that of simply decompressing and parsing the file (index creation induces ∼ 20% overhead) and with minimal working memory. The `mim` index itself is very small, usually about 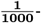th of the size of the original compressed file, and can be easily stored along side the file or fetched from a remote location when it is needed. Further, the `mim` index is purely additive — it does not modify the original `gzipped FASTQ` file in any way, nor require that the file be recompressed or rewritten — and thus it does not require converting the massive back catalog of existing raw sequencing data.

To demonstrate the feasibility and utility of the `mim` index, we benchmark construction of the `mim` index on a variety of existing `gzipped FASTQ` data, and also measure thread-scaling of `mim` index-assisted parallel `FASTQ` parsing on a simple parsing/ decompression-related task. We find that, for the one-time cost of index creation, and a small fraction of extra storage space, the `mim` index can massively accelerate the ingestion and parsing of `gzipped FASTQ` data, exhibiting near linear thread scaling in our experiments. `mim` is written in `C++`17, and is available as open source software under a BSD 3-clause license at https://github.com/COMBINE-lab/mim.

## 1. Introduction

Originally introduced around the year 2000, the text-based `FASTQ` format quickly gained adoption as the *lingua franca* of primary (“raw”) sequencing data distribution. It was subsequently adopted by tools, and now is the common format used by tools from *k*-mer counters ^1–3^, to read aligners ^4–7^, to assemblers ^8–10^, to transcript quantification tools ^11, 12^. The widespread adoption of the format has led to an inertia that has made it difficult to replace `FASTQ` with other formats that are more suitable to the massive scale of modern genomics data and the massively parallel character of modern computer architectures and algorithms. Moreover, due to the verbose and text-based nature of the `FASTQ` format, these files are often stored and shared in `gzip` compressed form. The European Nucleotide Archive (ENA) ^13^ currently contains approximately 63 Petabytes (PB) of data in the `fastq.gz` format.

While `gzip` compression drastically reduces the space required to store these files, and can help reduce the I/O burden between the storage and a running process, it imposes further efficiency constraints on how quickly the data can be processed. The fundamentally serial nature of the `gzip` format — the fact that, by construction, it is designed only to be read start-to-end ^14^ — stymies efforts at performing efficient parallel processing from raw data, making within-file parallel processing largely intractable.

### The bottleneck of parallel decompression and parsing

Combined with the fact that the development of new computational methods and pipelines leads to frequent re-processing of large collections of existing datasets, this means that a tremendous amount of time is wasted in decompressing and parsing data; for example, depending upon the hardware, lightweight tools for single-cell RNA-seq quantification^12^, or single-cell ATAC-seq quantification^15,16^ saturate in performance at 8-12 threads per input sample, yet continue to scale near linearly if the input is spread across multiple samples. Thus, this decompression and parsing task becomes a bottleneck when the tools being developed and adopted are increasingly fast and parallel. Some of these issues actually arise from the `FASTQ` format itself, but they are further exacerbated by dealing with `gzipped` versions of these files. Summarizing from Langmead et al.^17^, some of the main issues that limit parsing throughput include:

- Variable record lengths in `FASTQ` files can impede scaling because identifying these record boundaries needs to be done in a way that can maintain synchronization between files (e.g. in the case of paired-end reads);
- Synchronization and locking while reading a single `FASTQ` file by many threads can significantly affect performance; existing methods can approach this through an explicit locking mechanism between workers, or by using a single-producer multi-consumer approach where a dedicated thread decompresses and parses records — both limit scalability;
- `FASTQ` files are often stored in `gzipped` compressed formats which prevents efficiently reading these files using many threads, since decompression is a sequential process.

The paper of Langmead et al.^17^ also provides a good analysis of these bottlenecks for `FASTQ` and `gzipped FASTQ` files.

### Example

For example, it takes 104 minutes to compress a 132GB `FASTQ` file into 24GB `FASTQ`.gz, whereupon we loose the possibility of random access. Further, it takes 12 minutes to decompress and parse this file. In fact, it takes 9.5 minutes to simply decompress this file, without parsing, which, due to its sequential nature, can not be sped up using multiple threads without substantial overhead. This single-threaded decompression can be the main bottleneck in modern data-processing algorithms, which do benefit from multi-threading and parallelization, and often make use of SIMD instructions. Our index solves this by creating *checkpoints* every 32MB (by default), so that multiple threads can work on decompression and parsing in parallel without global locks. In our example, this speeds up the decompression and parsing from 12 minutes to just 37 seconds using 24 threads.

### Challenges with gzip

With continued advancements in sequencing, the amount of data being generated, and the corresponding sizes of the `FASTQ` files being produced is increasing. For example, Kerbiriou et al.^18^ show that `gzip` achieves a good size reduction for `FASTQ` files while still offering reasonably fast decompression. However, reading data from a compressed archive is still a relatively slow process, with *gunzip*, the decompressor component of *gzip*, yielding data at about 50-250 MB/sec, which is one to two orders of magnitude lower than the read throughput of current SATA/NVMe solid-state drives, depending on the technology.

The `gzip` application uses the `DEFLATE` algorithm^19, 20^ to compress and decompress binary data. `DEFLATE` consists of two stages. In the first stage, the data is processed sequentially and LZ77^21^ parsing is performed. This encodes the data as a sequence of literals and (off,length) pairs. (off,length) is the offset and length of the longest prefix in preceding 32KB context that can be used to replace data at this location. Once the LZ77 parsing is complete, Huffman coding^22^ is performed to further encode the data compactly. The Huffman code trees are reset every block to give best results. `DEFLATE` decompression is the opposite process — the compressed data is decoded by Huffman decoding. This writes down the data in a 32KB circular buffer, which is then used by LZ77 decoding to decompress the data entirely. The sequential nature of the compression and decompression strategy adopted by `DEFLATE` makes random access, or even parallel decompression, using compressed `FASTQ` files hard.

### Block compression

Block compression in this format is possible, whereupon files are first broken into chunks, and then these chunks are compressed separately. In fact, this approach was pioneered in the bioinformatics community with the development and adoption of the `BGZF` format^23^. This can allow parallel decompression of the compressed file, but it raises additional issues. First, such an approach can negatively affect the compression quality obtained by the compression algorithm (though, depending on the specific input and the quality of the compression chosen, this is not always the case practically). Second, even if blocks are compressed independently, there is no guarantee that they are “record aligned” (i.e. that chunks start and end on `FASTQ` record boundaries, or that the blocks in paired-end files are synchronized and each contain the same number of records). Finally, this approach requires the file to be compressed using this approach itself, which is true of *some* but certainly not all existing compressed `FASTQ` files. The solution we present here works equally well with `BGZF` compressed `FASTQ` files as with standard `gzip` compressed `FASTQ` files.

### Semantic checkpoints

The compression and decompression algorithms used in `gzip` are inherently serial. As such, existing `FASTQ` parsers can only begin reading starting from the beginning of each compressed `FASTQ` file. However, if we know the relevant context of the compressor (32KB of uncompressed data before the checkpoint of interest), then we can then start decompressing only from that point onward. This is the core idea behind our method. We create certain “checkpoints” in the file that are approximately evenly-spaced with respect to the *uncompressed* text. For each checkpoint, we store the information necessary to begin decompression directly at this checkpoint. Likewise, to give *semantic* meaning to these checkpoints, we also store the rank of the first read record that begins at or past this checkpoint, and the byte offset where this read starts beyond the checkpoint. Using this idea, we can have multiple threads read the compressed `FASTQ` file, where each thread starts processing from a checkpoint and reads some number of records. Moreover, we can jump to arbitrary checkpoints within multiple files, allowing us to perform a quick “synchronization” in the context of e.g. paired-end reads, whereby many threads can subsequently process the reads independently after this initial between-file synchronization.

### The mim index

We have developed `mim`, a tool that creates an index over `gzip` compressed `FASTQ` files, as well as `mimparser`, a corresponding (`C++`) `FASTQ` parsing library that uses this index to be able to read the compressed `FASTQ` file in parallel from multiple threads. The idea is to incur a one-time cost of building the index, which then significantly accelerates parsing compressed `FASTQ` files in the future. The `mim` index is per-file, and so index-creation can be trivially parallelized across datasets. Further, although not yet implemented in our library, it is easy to build the `mim` index as a byproduct of reading an existing `gzip` compressed `FASTQ` file. We envision that the `mim` index can be shared alongside the `FASTQ` files in repositories like the European Nucleotide Archive^13^, which amortizes the cost of the index creation. Moreover, as the index is purely “auxiliary”, it can be used in tools that take advantage of it without *any* negative backward compatibility consequences for existing tools that are unaware of this index. Finally, the index itself encodes a cryptographic (blake3)^24^ hash of the file on which it has been constructed, which enables uniquely binding an index to a source `FASTQ` file allowing, e.g., content-based systems transparently fetching these indices from a remote source. When running a simple task such as counting the number of occurrences of each nucleotide in a `fastq.gz` file, the `mim` index enables near-linear speedup with the number of used worker threads.

## 2. Related Work

There are several tools and methods related to `mim` and what is proposed here, but none of them achieve precisely the same aim or strike the same balance. First, there has been substantial effort put into parsers that are well-engineered and efficient, and several that take advantage of multiple threads when the input consists of multiple samples/files. Second, there are several tools that have been developed to enable parallel decompression of existing (un-modified) `gzipped` files. These approaches require no modification of the original file, nor do they require the construction of an index over the file. Finally, there have been proposals to replace (`gzipped`) `FASTQ` wholesale, with formats better designed for efficient parsing and for more modern architectures and algorithms.

### Efficient (gzipped) FASTQ Parsers

Before being processed, the gzipped `FASTQ` records must be decompressed and parsed, and considerable engineering optimization has been put into efficient libraries for this purpose. For example, the popular, single-threaded `kseq`^26^ library can be used for this purpose, and it provides a highly-efficient parser implementation, written in `C`, that can work transparently over either a compressed or uncompressed input. This library has been wrapped and made available in other languages such as Python. A fresh implementation of `kseq`, written in modern `C++` is available in the `kseq++`^27^ library.

Since decompression is inherently sequential, most multi-threaded libraries use a producer-consumer approach. For example, FQFeeder28 uses one thread per input file (or file pair `FASTQ` parsing) and uses `kseq++`^27^ for the underlying parser; multiple input sets can be decompressed in parallel, but the the degree of parallel decompression and parsing possible is limited by the number of samples (files or file pairs) being parsed. The producer threads decompress and parse the records, and place *batches* of parsed `FASTQ` records onto a concurrent queue. Consumer threads then take these batches of parsed records and perform the actual computation (e.g. alignment, quality assessment, etc.). A similar strategy is followed by `RabbitFX`^29^, which also has additional optimizations, as well as by Rust libraries seq_io^30^ and paraseq^31^.

While these parsers focus on efficiency, they lack the ability to decompress and parse input from individual `gzipped FASTQ` files in a highly parallel manner. With these approaches, the producer thread pool cannot meaningfully have more threads than the number of files being parsed (since each file is decompressed and parsed sequentially). We see that this as the major scalability bottleneck for pipelines relying on this approach, a bottleneck that our tool solves.

### Parallel gzip decompression algorithms

`pugz`^32^ is a parallel algorithm that enables fast decompression of `gzipped` files. It can achieve this by being able to decompress from a random offset by creating the fully decompressed 32KB context using a two-pass heuristic approach. The 32KB context can be constructed almost always at low compression levels, while some approximations are needed at higher levels. `rapidgzip`^14^ expands upon `pugz` by improving the heuristic used for speculative decoding, as well as by improving the parallelization and cache efficiency of the underlying algorithm. Both `pugz` and `rapidgzip` can substantially improve the decompression speed of `gzipped` files, including `gzipped FASTQ` file. However, this improvement in decompression speed comes at a considerable cost of extra computation; as a non-trivial amount of the speculative decoding ends up in wasted work. The `rapidgzip` tool also exposes an “index” mode, whereby an extra index aids in parallel decompression of the file. While this approach is similar to our approach (and to that of the block `gzip` format with an index), it differs in that `rapidgzip` is a generic method, and therefore lacks a semantically aware index like `mim` which both enables synchronized decompression between related files (e.g. ends of paired-end reads) and allows lock-free and wait-free parsing of the underlying content, since `mim` checkpoints are, by construction, aligned to record boundaries.

### Modified gzip archives

`mgzip` and `pgzip` are Python libraries built on Python’s `zlib` and `gzip` wrappers. `pgzip`^33^, is a maintained fork of `mgzip`^34^, and compresses uncompressed buffers by breaking them into blocks, and compressing each block independently in parallel. The metadata of all blocks is inserted into the `FEXTRA` field within the `gzip` file. This allows for parallel decompression by reading this metadata. While they maintain backward compatibility with `gzip` (i.e. the `gzip` utility can decompress the generated `gzip` file), they generate a different `gzip` file due to independent block compression. Thus, adopting such an approach requires re-compressing the data in this specialized format. Likewise, as with the index adopted by `rapidgzip`, the blocks of `pgzip` and `mzip` are not aware of the semantics of the underlying file, and therefore not necessarily aligned with records.

A related approach is that taken by the Blocked GNU Zip Format (`BGZF`)^35^. In this format, small chunks of the original file (≤ 64KB) are compressed independently as blocks with a fresh compressor state for each block. This allows the blocks to be decompressed in parallel. The `BGZF` format essentially represents a multi-archive `gzip` file where the independently compressed blocks are concatenated together (which constitutes a valid `gzip` file). The header of each `BGZF` block also contains extra information about the length of the compressed block. The `BGZF` format is also designed to support an index, which is a simple list of offset pairs specifying the position in the compressed stream of the start of each block, as well as the uncompressed position to which it corresponds, this makes it easy to jump to and decompress individual blocks, as well as to perform random access within the compressed file. While the `BGZF` format (with default parameters) does make an effort to be somewhat semantically aware of the underlying data being encoded (i.e. the specification states that “`bgzip` will attempt to ensure `BGZF` blocks end on a newline when the input is a text file.”), it still lacks specific information about the ranks of the underlying reads. Moreover, there is no guarantee that a newline boundary corresponds to the start or end of an entire read record (e.g. `FASTQ` records typically span 4 lines, and (non-multi-line) `FASTA` records span 2). This, therefore, makes it impossible to properly align multiple `BGZF`-compressed files and to process them in parallel in a synchronized fashion (as is necessary e.g. with paired-end reads). Likewise, while the `BGZF` index is very simple and also very small, it relies on this special structure of `BGZF`, and so it is not applicable on general `gzip` compressed `FASTQ` files, as is `mim`.

### Alternative storage formats

While the approaches above all focus on either maintaining existing `gzip` archives as-is, or generating special variants of `gzip` archives that are more amenable to random access or parallel decompression, researchers have also considered what might be possible if the `gzip` and `FASTQ` formats themselves could be replaced as a source of raw data storage and transfer. For example, the Nucleotide Archival Format (NAF)^36^, describes a format designed to be smaller and simultaneously faster to decompress than `gzipped FASTQ` files.

Recently, the Binseq^37^ family of formats (comprising `BQ` and `VBQ`) was proposed. These formats specifically focus on encoding high-throughput sequencing data in a manner that can be processed in parallel (i.e. either by enforcing fixed-size records in `BQ` format, or by adopting a record-aligned block/chunk compressed structure in `VBQ` format). These approaches enable, by design, massively parallel decompression and parsing, and can achieve precisely the types of speedups we hope to enable. Yet, for the NAF format and the Binseq format family, as well as for related alternative approaches, the main impediments are twofold. First, these formats are not well-represented in the massive existing repository of sequencing data, and the conversion of all prior data into these more efficient formats would be a massive undertaking. Likewise, tremendous care would have to be taken to ensure that no important aspects of the existing data were lost in the process, for example, due to unexpected errors during conversion or potential bugs or corner cases in the encoder for the new format. With the purely auxiliary `mim` index, the original file is only ever read (not rewritten or modified), and so even a failure to index runs no risk of losing information from the original file. Second, even if this conversion was undertaken successfully, the vast majority of existing bioinformatics tools do not support these formats yet (and many never will), and so, to retain the utility of the vast repository of existing data, one would likely still have to repeatedly convert from these newer formats back into `FASTQ` or `gzipped FASTQ` when processing with legacy tools. This burden may be minimized if the conversion itself is lightweight and fast, *and* if the downstream tool being targeted is capable of consuming input from a FIFO or named pipe. On the other hand, the approach we propose maintains full backward compatibility by default (the source files are unchanged, and the `mim` index is purely auxiliary). Thus, no large-scale conversion need be performed, the existing repository of available data can be retained as it currently exists, and no intermediate conversion is necessary to allow data to be ingested by existing tools. New tools can choose to take advantage of the auxiliary index we propose in an iterative fashion, and legacy tools can be retrofitted, as desired, to do so.

## 3. Our Approach

### 3.1. The mim index

In this section we describe the design for `mim`, the index structure we use to enable parallel parsing of `gzipped FASTQ` files.

#### 3.1.1. The zran index

The `mim` index builds upon the `zran` index. `zran` is an example application that appears in the source tree of the `zlib`^38^ library. `zran` demonstrates the use of certain `zlib`^38^ API features to enable accessing a `gzip`-compressed file from a random offset.

It accomplishes this by fully decompressing the file, and building an index containing “access points” (which we henceforth call checkpoints) at approximately equally-spaced locations, as shown in Figure 2. Each checkpoint includes the starting file offset, relevant metadata about the current deflate block, and the necessary data to exactly reproduce the state of the decoder at this checkpoint (i.e. the decoder’s context). This index is then used to fetch some number of bytes from an arbitrary, user-provided location in the original `gzip` file (starting at the most-recent checkpoint prior to the desired location). This program does not save or load the index from disk, but rather stores it in memory for a one-time random fetch (`zran` is, after all, an example and not a full application).

**Figure 1.**
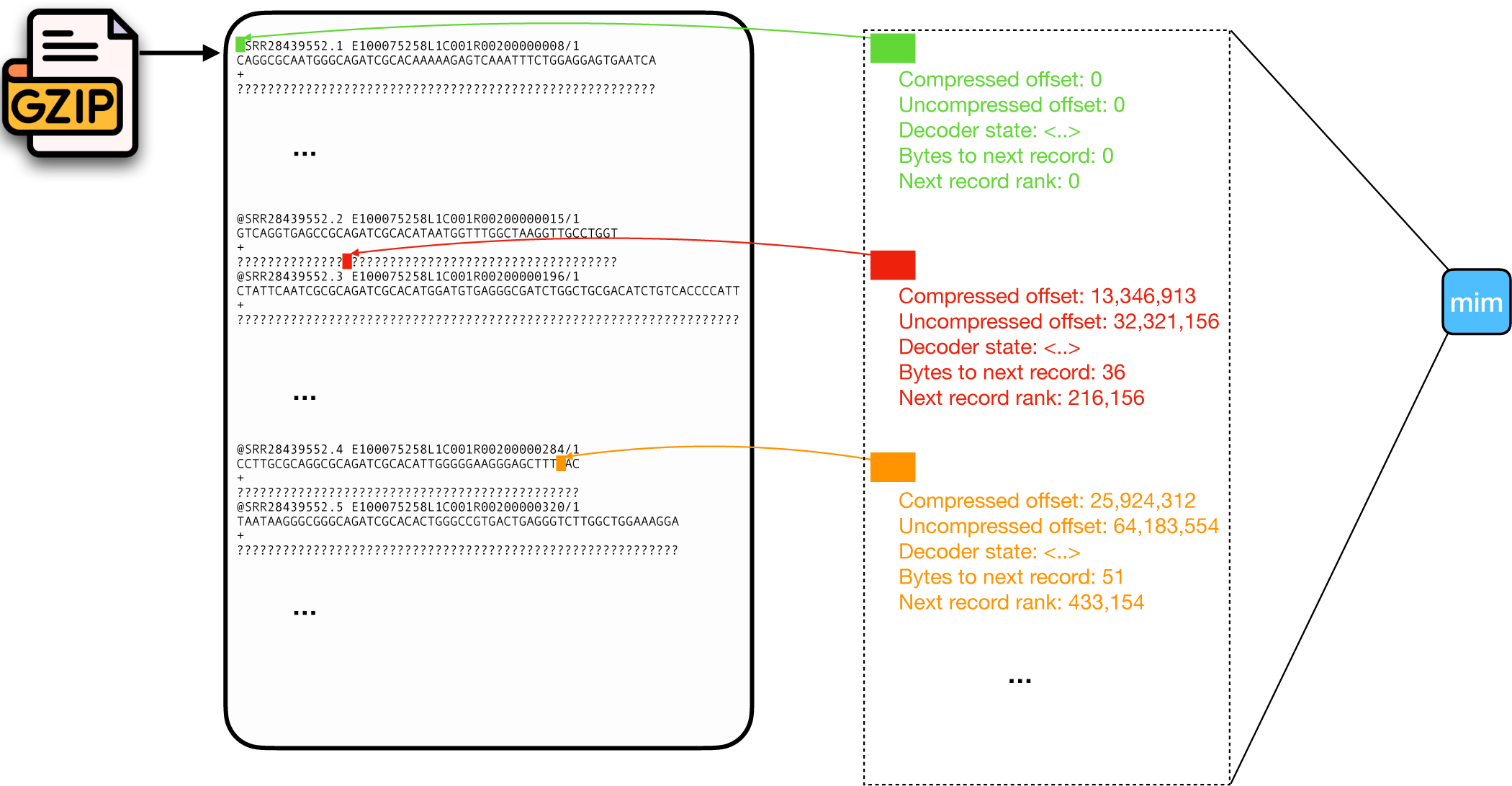
Overview of the structure of the `mim` index. Several checkpoints are shown, along with the important information they retain. The logo for Gzip is taken from^25^.

**Figure 2.**
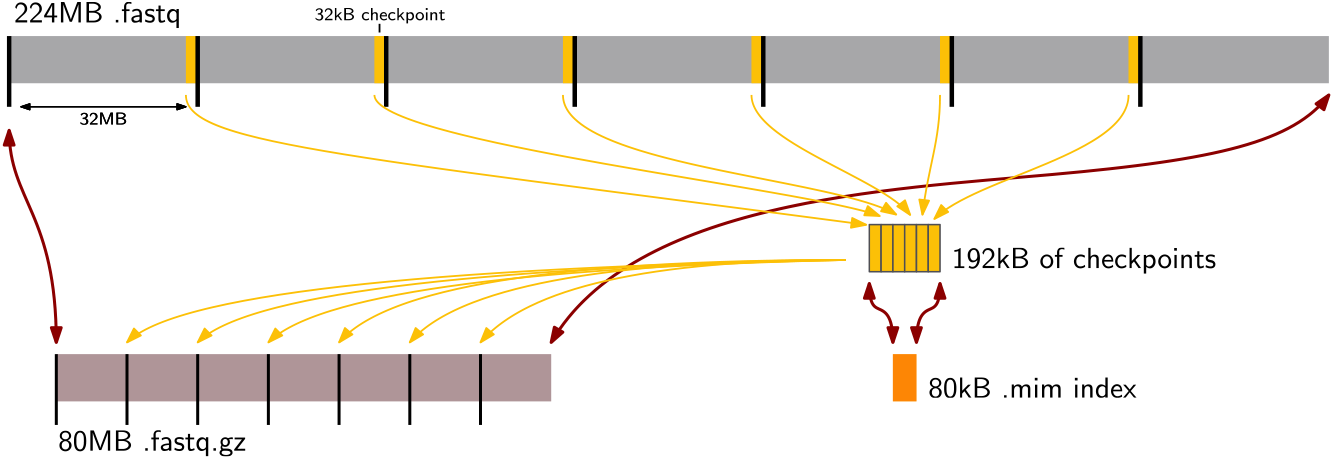
Schematic example of the mim index. A 224MB `FASTQ` file (top, grey) supports random access, but its 80MB gzipped `fastq.gz` representation (bottom left, dark red) does not. The `mim` index makes a checkpoint (black bars) after approximately every 32MB of plain text. For each checkpoint, it stores the internal state of the gzip decompressor, which consists mostly of the last (up to) 32kB of sequence (vertical yellow bars, not to scale), and some small metadata (not shown). The data for all checkpoints is concatenated with the metadata and gzipped to obtain the `mim` index file (bottom right, bright red). To read the `fastq.gz` file in parallel, the `mim` index is first decompressed to obtain the checkpoint data. Then, each thread is assigned a range of 32MB blocks and can decompresses and parse this range by initializing the decompressor with the checkpoint data.

#### 3.1.2. Augmenting zran for sequencing data

`FASTQ` and `FASTA` files contain records that are not of fixed size, and so each record can span a variable number of bytes. The `mim` index and `mim-parser` handle both `FASTQ` and `FASTA` files, but here we will focus only on `FASTQ` for ease of exposition.

Any application that reads `gzipped FASTQ` files efficiently in parallel needs to be able to (1) distribute work over multiple threads (2) move the parsing of records outside of any critical section of the program and (3) ensure that the work assigned to different workers can be decompressed efficiently in parallel. To achieve these capabilities, the `mim` index augments `zran`’s index to support record-aligned access.

We achieve this by maintaining, for each `zran` checkpoint, two additional pieces of metadata; (1) the byte offset of the first record occurring at or after this checkpoint, i.e. how far from the checkpoint, in the *uncompressed* stream, one must read to encounter the start of the first complete record in this chunk, and (2) the record rank corresponding to this record, i.e. starting from 0, how many records have we seen prior to this one in the uncompressed stream. To compute this auxiliary information, we augmented the `kseq++`^27^ parser to store, for each parsed record, the byte offset from the start of the file until the record start position. This “semantic” mapping, that puts `zran` checkpoints into correspondence with specific reads and stores check-point-local offset from the start of the next record, is built directly after the original `zran` index is built, and is stored alongside the standard `zran` index as a list of checkpoints. Figure 1 shows an overview of the `mim` index.

We store, along with our index, a cryptographic hash of the content of the *compressed* file. Specifically, we store the `blake3`^24^ hash of the compressed file content. We envision this data is being useful for two reasons. First, we hope, in the future, to be able to host `mim` indexes for a wide variety of publicly available datasets online. Having a distinct content-based key will make it easy for `mim` enabled parsers to automatically and transparently download and use the `mim` index for a file if it is available (and if the user wants to permit this behavior). Second, this provides a failsafe against accidentally using the wrong `mim` index with a `gzip` file to which it does not correspond. While attempting to parse a `gzipped FASTQ` file in parallel using a mismatched `mim` index will anyway likely lead to a failure to parse, the checksum adds an extra level of certainty.

Finally, the index can, upon creation, embed user-provided metadata. This metadata is provided as a `JSON`^39^ object (serialized in the index header in `CBOR`^40^ format). This allows the user to tag the index with useful descriptors, relevant provenance information, or other details that should be linked to the associated file.

#### 3.1.3. Aspects of the mim index

In order for the index to be useful, the compressed file needs to have a sufficient number of checkpoints to saturate many parsing threads. Specifically, we want each chunk not to be *too* large, as, when we need to synchronize records between files (e.g. for the case of paired-end parsing), we may need, in the worst case, to decompress and discard up to one chunk’s worth of data in each thread. Of course, having more chunks and more checkpoints makes the index itself larger on disk, though we mitigate this somewhat by compressing the index itself on disk when we store it. When building the `mim` index, the *span* (average number of uncompressed bytes between stored checkpoints), is a user-controllable parameter that determines the number of checkpoints that will be stored. We set a default span of 32MB, and that is the span size used for all data presented in this manuscript.

Ultimately, we observe the size of the index to be very small; ∼ 0.1% of the size of the compressed data file itself in most cases, since each checkpoint is around 32KiB (∼ 32MB/1000, despite the change in base). In cases where the input file is actually already a `BGZF` file (i.e. was `bgzip` compressed), we find the index to be even smaller (0.0003% of the size of the file for `pbmc_10k_v3_S1_L001_R2_001`, or only about 96B per checkpoint). This is because `zran` chooses its checkpoints naturally to align with the `bgzip` chunks (each of which is typically much smaller than the chunk size in our index). Since each `bgzip` chunk requires no additional context to decompress, our index collapses to store basically just the auxiliary chunk to read offset and read rank information.

Index creation itself requires decompressing and parsing the original input file, plus a small amount of bookkeeping to construct the `zran` and auxiliary index itself. Thus, the time to create the `mim` index is only about ∼ 20% larger than the time required to simply parse the `gzip`- compressed `FASTQ` file. We note here that we have not made an attempt to optimize the `mim` index creation, and it can, almost certainly, be done in a single pass over the input file. Further, this index creation cost is is a one time cost. Therefore, we believe that this is a reasonable tradeoff. Once created, the indices themselves are very small (the largest we encountered in our testing was ∼ 25MB, and load within a fraction of a second. Statistics concerning the index sizes and how they relate to the original files are given in Table 1.

**Table 1.** The size of each dataset evaluated and the size of the corresponding mim index and the relative size of the mim index as a percentage of the original file. The pbmc file, designated with a *, was block compressed using bgzip.

| dataset (.fastq.gz) | size (GB) | # of checkpoints | mim index size (MB) | size reduction |
| --- | --- | --- | --- | --- |
| SRR28048028 | 0.390 | 74 | 0.407 | 958× |
| SRR28028283_1 | 2.47 | 617 | 2.59 | 954× |
| SRR28028283_2 | 2.51 | 617 | 2.63 | 952× |
| SRR28439552_1 | 3.62 | 704 | 3.74 | 967× |
| ERR1190770_2 | 4.06 | 346 | 4.19 | 970× |
| SRR9331208_1 | 7.93 | 992 | 8.13 | 974× |
| SRR13060943_2 | 8.56 | 907 | 8.80 | 972× |
| ENCFF000FFF | 8.84 | 869 | 9.10 | 971× |
| SRR26865528_2 | 18.4 | 3902 | 19.2 | 959× |
| SRR28896749_1 | 24.7 | 4428 | 25.5 | 965× |
| pbmc_10k_v3_S1_L001_R2_001* | 16.7 | 2420 | 0.0473 | 353 154× |

### 3.2. mim-parser

Next, we describe our design for the parallel parser that reads `gzipped FASTQ` files taking advantage of the `mim` index. One major benefit of the approach described above is that obtaining disjoint and independent, record-aligned chunks of the input file, requires almost no synchronization. In our parser, upon loading the initial index, the parser evaluates the number of chunks in the input file in light of the number of parallel worker threads that have been registered. It then assigns contiguous intervals of chunks to the worker threads such that the number of chunks handled by each worker is as even as possible. After this initial assignment of chunk intervals, *no synchronization* is required between the worker threads. Each consumer/worker thread retains a read-only reference to the `mim` index, it jumps to the beginning of the chunk interval it was assigned, and determines the number of records that it is requested to process. This value is obtained by subtracting the rank of the first read it is assigned (i.e. the read at the start of its first assigned chunk) from the rank of the first read in the chunk after its assigned interval. Then, each worker simply begins reading and parsing the input file from its assigned location, and yielding records until it has produced the requested number of records.

The records that are read are parsed using `kseq++`^27^ and converted into a `C++` structure that can be used by downstream tasks. Specifically, we have made a wrapper for `kseq++`’s input stream type to allow it to read directly from a particular offset in a `gzip` file where the decompressor state has been properly primed by an index checkpoint. This is possible because the `kseq++` parser provides a user-implementable input stream interface that need only implement the ability to obtain a requested number of bytes of uncompressed data from the input stream. Specifically, we open the `gzip` file, seek to the appropriate compressed offset, and prime the decompressor with the necessary dictionary state. Then, we determine, based on the record-level information associated with this check-point, how many bytes we must discard before the first read record occurs. We read and discard this number of bytes, and subsequently the file can be read by this worker thread, from the start of a record, directly as if it is reading from a normal `gzip` compressed stream (our stream reading implementation automatically handles appropriately skipping `gzip` headers in a `BGZF` file or in a multi-part `gzip` archive).

The parsing of paired-end read files proceeds in a similar manner but is slightly more sophisticated. Here, the challenge is that the parsing of records between the two files must be synchronized so that, e.g. the rank *k* read from file 1 is parsed and returned along with the rank *k* read from file 2. To achieve this, the indices for both files are opened, and the intervals of chunks are assigned to the worker threads, based on the chunks in read file 1, just as described above. Now, each worker knows what range of reads it will parse and produce from file 1. Let *r* be the rank of the first read that will be parser from file 1 by worker *w*. Next, for this worker, a search is conducted over the checkpoints in the index for file 2 to find the chunk starting with the highest ranking read ≤ *r*. Let this chunk in file 2 be called *c*, and let the rank of the first read in this chunk be *r*^′^. The worker then opens the second `gzip` file starting at chunk *c*, discards the bytes prior to the start of read *r*^′^, and then parses and discards *r* − *r*^′^ read records. At this point, the worker is situated at the same rank read in files 1 and 2, and it can simply process the records from these files in sequence (and independent of all other workers) until it has processed the prescribed number of reads. This initial synchronization means that some small number of records may be read and parsed more than once (e.g. discarded by one worker to obtain synchronization, while properly yielded by another as part of its assigned work interval). Nonetheless, this constitutes only a small amount of extra work once per thread (per file), and in practice represents negligeable overhead. We note that it *would* be possible to design a `mim` index directly for paired-end reads that records the exact points and offsets in the file pair for reads of matching ranks, eliminating this small amount of extra work. However, we decided against such an approach, as we find the current approach simpler. For example, the process of index creation is independent for each compressed file, and a file can be indexed (and parsed) independently of the file with which it is paired. Likewise, The approach we adopt can easily and naturally be extended to file sets of arbitrary arity using their independently computed indices (e.g. to single-cell ATAC-seq data which might require sets of 3 files that are parsed in a synchronized manner).

## 4. Evaluation

In this section, we discuss our experimental setup and the results for benchmarking `mim` and `mim-parser`. All experiments were performed on a single server with two Intel Xeon E5-2699 v4 2.20 GHz CPUs having 44 cores, 512 GB of 2.40 GHz DDR4 RAM, and a number of 3.6 TB Toshiba MG03ACA4 ATA HDDs. The system is running with Ubuntu 20.04 GNU/Linux 5.4.0-172-generic. The running times are measured with the GNU `time` command. To collect the running times, the initial single-threaded parsing was performed first, and the command was issued twice recording only the timing of the second run, to ensure a warm cache. Subsequently, the times were recorded for the same sample for the remaining thread counts.

### 4.1. Experimental setup

We seek to evaluate our approach with respect to two critical criteria: (a) Evaluate how lightweight the `mim` index is, and (b) determine how much speedup can be achieved via this kind of parallelism.

For the first aspect, we evaluate the index creation times and index sizes for a range of `FASTQ` files of different sizes (and characteristics).

For the evaluation, we kept the span of checkpoints to be constant (every 32, 000, 000 uncompressed bytes in the underlying stream), which generated different index sizes for different `FASTQ` files. Ideally, we may want to optimize checkpoint placement for a given file size or file structure – as for large files, this span between checkpoints generates *many* more checkpoints than we need to obtain maximal parallelism — but we leave this for future work. Nonetheless, this sampling rate between checkpoints leads to indices that are about 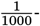-th the size of the original file, with little variance observed over these datasets.

To evaluate the speedups we can achieve with `mim-parser`, we run `mim-parser` on 11 different `FASTQ` files (including 2 files that constitute a paired-end dataset, which we process both individually and as a synchronized pair), with details given in Table 1. These data represent a collection of different assay types (e.g. metagenomic sequencing, ChIPseq, RNA-seq), read lengths, data ages and file sizes, with sizes ranging from ∼ 400MB (compressed) to ∼ 24GB (compressed). We run the parser in each case varying the number of threads from 2 through 24.

To keep the focus on the speed of *parsing* and not any computation done with the parsed data, the downstream task used for benchmarking is a simple task of counting the number of occurrences of each nucleotide across all of the records in each of the `FASTQ` files. We compare our tool’s performance with a sequential reader using the same underlying parsing library.

### 4.2. Results

#### 4.2.1. Evaluating mim index

Table 1 shows the index size for `FASTQ` files of different sizes. The index size is about 0.1% of the size of the original (compressed) `FASTQ` file. Of course, this index size can be controlled by changing the desired number of bytes between checkpoints (for all experiments here, we use 32, 000, 000 bytes between checkpoints).

In Table 2, we also report index creation times. On average, index creation takes ∼ 1.2 × as long as simply parsing the file with a single thread. This is expected, as index creation is (currently) a single-threaded process that has to stream over the file twice; once, without any parsing to generate the `zran` checkpoints, and again with parsing to generate the semantic information for each checkpoint. Optimization of the index creation step is one concrete direction for future work. However, we believe that the current index creation speed is already acceptable, as it is a one-time cost and will be amortized over many instances of processing the resulting files. Further, since index creation is completely independent per file (even for files that are paired in sequencing), the process of index creation can be trivially parallelized over separate files.

**Table 2.**
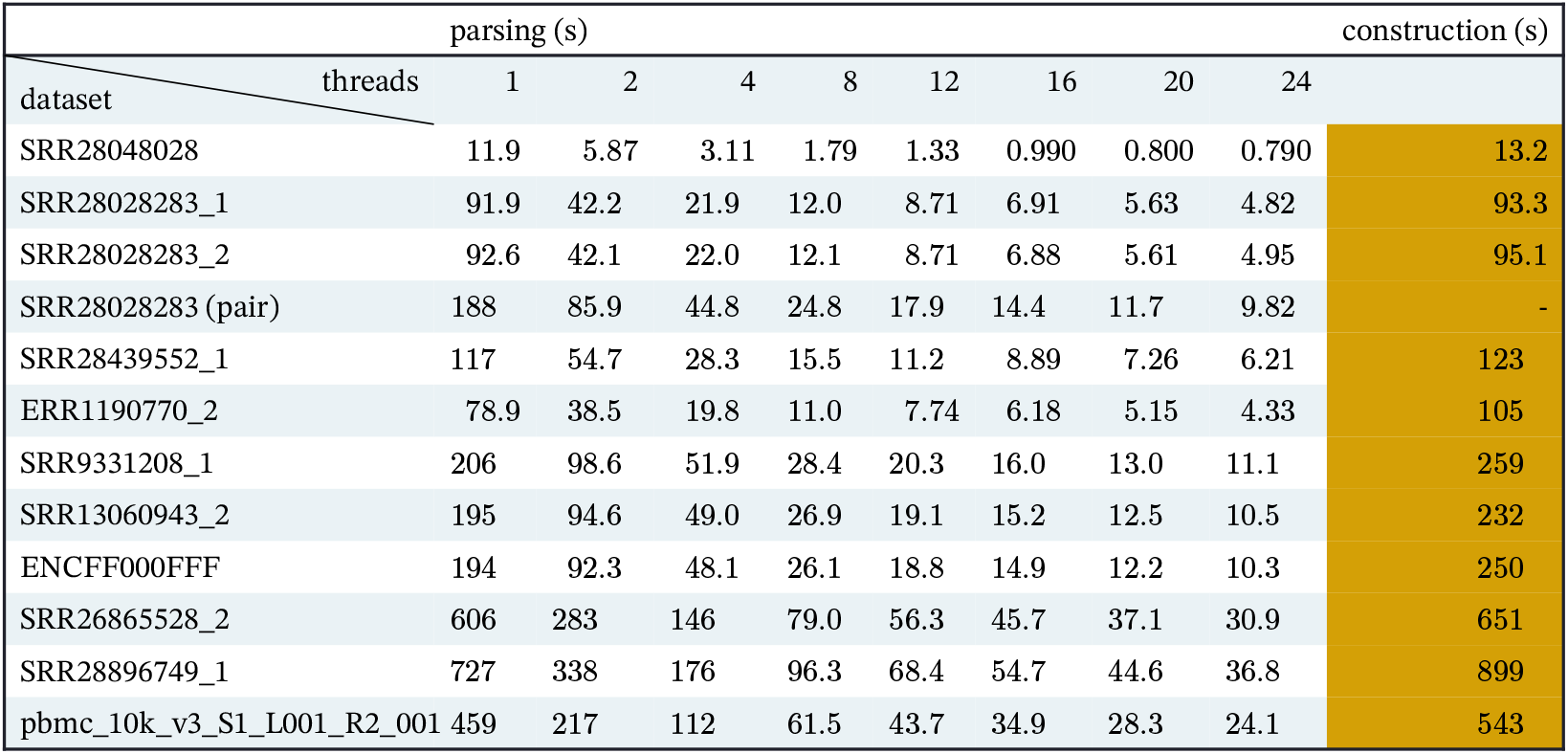
Time required to construct the mim index and to compute the nucleotide sums using the mim-parser with varying numbers of threads.

#### 4.2.2. Evaluating mim-parser

Figure 3 shows the scaling results for `mim-parser` on the set of `FASTQ` files listed in Table 1. We can see that `mim-parser` scales near linearly with the number of worker threads for these files, at least up through about 8 threads, where we likely start to hit the limits of the HDDs being used in these experiments.

**Figure 3.**
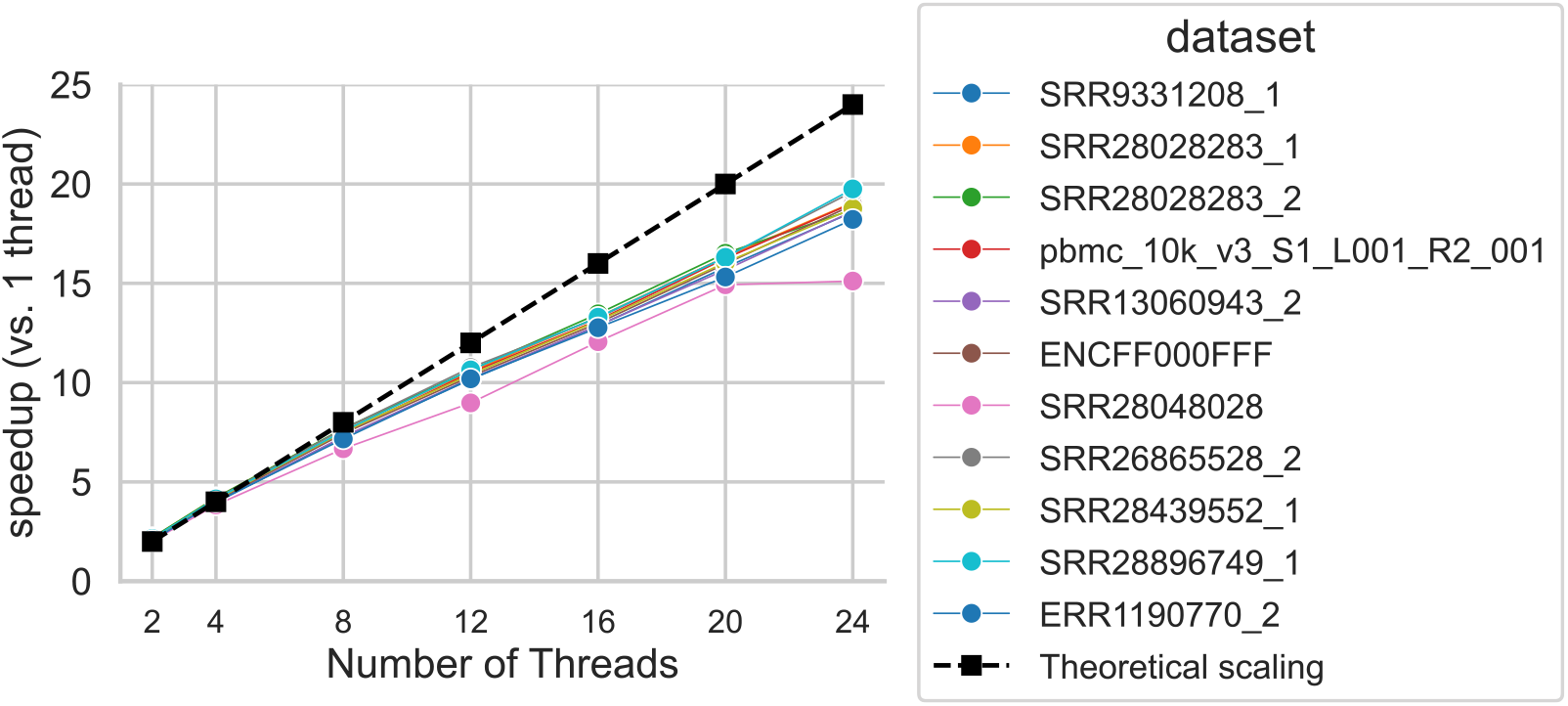
Scaling results for `mim-parser`. The plot shows the speedup of processing with *t* threads compared to 1 thread for each file while varying *t* from 2 to 24. The plotted value is simply the time to process that file taken by 1 thread divided by the time take with *t* threads. The black dashed line represents theoretically optimal scaling, where the speedup is simply *t*. Note that `SRR2848028` is the smallest file, and, though it has more than 24 checkpoints, the total execution time with 24 threads is only 0.79 seconds, so speedup is limited by other factors like index reading, initial work assignment, etc.).

## 5. Future Work

One substantial potential area for future optimizations is determining an optimal policy for the size of `mim` index (i.e. how to place checkpoints). Since the `mim` index builds on `zran`’s index construction algorithm, we currently create checkpoints that are separated by approximately the span requested by the user. However, once one has substantially more checkpoints than parsing threads that will ever be used (e.g. when one has at least a few hundred check-points), the creation of further checkpoints is likely of diminishing utility (though it may provide some benefit in speeding up the initial synchronization in paired-end parsing). Thus, a policy that determines how to place “enough” checkpoints, and where they should be placed, has the potential to make the `mim` index even smaller. We also store a mapping of record boundaries to their respective byte offsets in the `gzipped` file. One potential optimization that may be possible is to construct check-points exactly at record boundaries, so that these two data structures can be represented as one, helping us further reduce the size of the index.

Another area for potential improvement is a more efficient construction of the `mim` index itself. While the current construction takes only small memory and is linear in the size of the file being indexed, the index should, in theory, be possible to construct in a single pass. Moreover parts of the construction (e.g. the computation of the `blake3` hash) may even be possible to parallelize. We believe that the index is efficient enough to begin constructing at scale now for the vast catalog of existing data, but further optimization of the construction algorithm will certainly help reduce the time and financial resources required to perform this indexing.

Independent of index construction, our current parsing strategy breaks the compressed file into approximately equal intervals of data, and has each worker thread process its own interval independently. While this is very likely to work well in the vast majority of cases, one might imagine scenarios where equal quantities of input reads result in an unequal distribution of work in a downstream task (e.g. perhaps the user is performing alignment and some region of a file is enriched for reads that align to repetitive parts of the reference sequence). Alternative load balancing strategies are possible. For example, instead of each thread being assigned an approximately equal region of the file such that all regions cover the entire input, each thread could be assigned a disjoint region of the file such that the sum of all regions (all of which still start at a checkpoint) are much smaller than the total file length. In this case, when a thread is done with its assigned region, it can consult the parser for the next available region, etc. As the regions are made smaller, the work is broken up in a more fine-grained way, and the strategy naturally adapts to ensure that worker threads are not starved. Such a strategy requires marginally more coordination, but that is likely to be negligeable.

Perhaps the most impactful line of immediate work will be to begin computing and hosting `mim` indexes for publicly deposited sequencing data. While we can begin this process on a small scale using resources available to us, it would be ideal to run `mim` index creation in a massively parallel manner in the cloud, akin to how, e.g. the Logan project41 made use of AWS to perform unitig and contig assembly of the entire SRA^42^. Along with this computation of the `mim` index over the existing catalog of sequencing data, it will also be useful to provide a remote hosting of these indices so that they can easily be obtained given just the accession number of the corresponding read files. Even better, given the content-based cryptographic hash embedded in each `mim` index (i.e. the `blake3` hash computed from the compressed representation of the data), we would like to add to our parser library the optional ability to transparently download and use a `mim` index for a `FASTQ` file if one is available. The unique content addressability of the index based on the file makes this possible.

Finally, the current `mim-parser` implementation is written in `C++17` and usable by any tool capable of adopting this language. This is a reasonable choice for a first implementation given the prevalence of `C++` in the development of tools for high-throughput genomics processing. However, an increasing number of new high-performance genomics tools are being written in `Rust`, and we would like to provide a `Rust` implementation of a `mim-parser` (or to provide the requisite functionality as a crate that can be used by existing `Rust FASTQ` parsers). Likewise, we would also like to develop and expose `Python` bindings for the `Rust` implementation, to provide even broader support for using the `mim` index.

## 6. Conclusion

This work addresses a substantial bottleneck faced by many tools that need to process and parse raw sequence genomics data. We present the `mim` index, a lightweight, supplementary index that accompanies a `gzipped FASTQ` file and enables truly parallel decompression and parsing of the file by a large number of threads.

To demonstrate the utility of this index, we also provide an implementation of `mim-parser`, a multi-threaded `FASTQ` parser that achieves a near-linear speedup compared to serial parsing. By using the `mim` index, each of the workers in `mim-parser` can parse disjoint ranges of records within a compressed `FASTQ` file in parallel, and provide them for downstream processing. Our results show that `mim-parser` achieves strong scaling as we increase the number of workers. Moreover, the structure of the `mim` index allows the parallel decompression and parsing even of synchronized (e.g. paired-end) files, and this is also supported by `mim-parser`.

A key benefit of the `mim` index is that it is purely additive / auxiliary. That is, it does not require modifying or recompression of the source `FASTQ` files in any way. This means that adoption can be performed incrementally, and that pipelines can freely mix tools that support `mim` index enhanced parsing with those that don’t. Further, the indexes are small, easy to store and transfer, and have a strong built-in mechanism for correctness validation. Despite its relatively simple strategy, our proposed approach enables much better utilization of available modern hardware in the processing of raw sequencing data, and can lead to substantial performance gains for existing tools and algorithms that are currently bottlenecked on I/O and parsing. When a sufficient number of threads are available, this includes some of the most ubiquitous and common processing, like read alignment^17^. In summary, we hope that researchers find the `mim` index and `mim-parser` useful in speeding up their workflows, cutting costs, making better use of modern parallel hardware, and potentially making prohibitive tasks more accessible.

## Acknowledgements

The authors wish to thank Noam Teyssier, and Bede Constantinides for important conversations during the development of the `mim` index and `mim-parser` parsing strategies. The preliminary work that led to this project was completed as a semester project for the class CMSC701 at the University of Maryland, and Siddhant Bharti, Prajwal Singhania, and Rakrish Dhakal wish to acknowledge the Zaratan HPC cluster^43^ as an important resource in carrying out that work. Finally, Rob Patro wishes to thank Robert Aboukhalil, Robert A. Petit III, and Wytamma Wirth for motivating him, during an animated discussion at the CZI Open Science meeting, to pick this project back up after a prolonged hiatus and finish it.

## Declaration

R.P. is a co-founder of Ocean Genomics Inc.

